# Roadmap For The Expression Of Canonical and Extended Endocannabinoid System Receptors and Proteins in Peripheral Organs of Preclinical Animal Models

**DOI:** 10.1101/2023.06.10.544455

**Authors:** JJ Rosado-Franco, AL Ellison, CJ White, AS Price, CF Moore, RE Williams, LB Fridman, EM Weerts, DW Williams

**Affiliations:** Department of Molecular and Comparative Pathobiology, Johns Hopkins University-School of Medicine, Baltimore, Maryland, USA; Department of Microbiology and Molecular Immunology, Johns Hopkins University-Bloomberg School of Public Health, Baltimore, Maryland, USA; Department of Psychiatry and Behavioral Sciences, Johns Hopkins University Bayview Campus, Baltimore, Maryland, USA; Department of Neuroscience, Johns Hopkins University-School of Medicine, Baltimore, Maryland, USA; Division of Clinical Pharmacology, Johns Hopkins University-School of Medicine, Baltimore, Maryland, USA

**Author notes:** Correspondent author: Dionna W. Williams, Ph.D., Department of Molecular and Comparative Pathobiology 733 North Broadway, MRB 831, Baltimore, MD, USA 21205. Authors contributed equally (JJ Rosado-Franco, AL Ellison).

**Keywords:** Endocannabinoid system, receptors, cannabinoids, mice, rat, non-human primate

## Abstract

The endocannabinoid system is widely expressed throughout the body and is comprised of receptors, ligands, and enzymes that maintain metabolic, immune, and reproductive homeostasis. Increasing interest in the endocannabinoid system has arisen due to these physiologic roles, policy changes leading to more widespread recreational use, and the therapeutic potential of *Cannabis* and phytocannabinoids. Rodents have been the primary preclinical model of focus due to their relative low cost, short gestational period, genetic manipulation strategies, and gold-standard behavioral tests. However, the potential for lack of clinical translation to non-human primates and humans is high as cross-species comparisons of the endocannabinoid system has not been evaluated. To bridge this gap in knowledge, we evaluate the relative gene expression of 14 canonical and extended endocannabinoid receptors in seven peripheral organs of C57/BL6 mice, Sprague-Dawley rats, and non-human primate rhesus macaques. Notably, we identify species- and organ-specific heterogeneity in endocannabinoid receptor distribution where there is surprisingly limited overlap among the preclinical models. Importantly, we determined there were only five receptors (CB2, GPR18, GPR55, TRPV2, and FAAH) that had identical expression patterns in mice, rats, and rhesus macaques. Our findings demonstrate a critical, yet previously unappreciated, contributor to challenges of rigor and reproducibility in the cannabinoid field, which has profound implications in hampering progress in understanding the complexity of the endocannabinoid system and development of cannabinoid-based therapies.

## Introduction

The endocannabinoid system (ECS) has been evolutionarily conserved to preserve its importance in maintaining immune, metabolic, and reproductive homeostasis (1–4). This system is present in all vertebrate animals, including rodents, non-human primates (NHP), and humans (4,5). The canonical ECS is comprised of two main cannabinoid receptors (coded by the *cnr1* and *cnr2* genes), endogenous lipid ligands (endocannabinoids, i.e., anandamide and 2-arachydonoil glycerol), and enzymes involved in endocannabinoid metabolism (coded by the *faah* and *naaa* genes, among others not included in this study) (1,6). There are additional extensions to the canonical ECS, termed the “extended” ECS, that are comprised of receptors with primary functions in other pathways that have accessory functions that exist upon interaction with cannabinoids (7,8). Some of these receptors include peroxisome proliferator activated receptors (coded by the *ppara* and *pparg* genes, respectively), “endocannabinoid-like” G-protein coupled receptors (i.e., *gpr18*, *gpr55,* and *gpr119*), nociception ion channels (coded by the *trpv1* and *trpv2* genes, respectively), and transporters (i.e., *htr1a*, *adora2a,* and *adgrf1*) (9,10). Though their primary functions are best characterized in other pathways, the extended ECS receptors functionally interact with endocannabinoid ligands, the phytocannabinoids present in the *Cannabis* plant, and other endogenous lipid mediators, including oleoyl-ethanolamide (OEA), palmitoyl-ethanolamide (PEA), and linoleoyl-ethanolamide (LEA) (9,10). Together, the canonical and extended ECS, known as the “endocannabinoidome”, consists of many receptors that can interact with multiple ligands, thus creating a complicated network of outcomes during both health and disease and not limited to the brain.

More widespread accessibility of phytocannabinoids for medicinal and recreational use, policy changes that have impacted funding priorities, and the heightened desire for plant-based therapeutics have re-awakened scientific interest in the ECS. As such, preclinical animal models are becoming increasingly important in identifying the health implications of phytocannabinoids and the molecular mechanisms by which the ECS can be therapeutically leveraged. However, challenges exist in the translational capacity of preclinical studies due to conflicting reports that arise because of key differences in study design, including the route of administration, formulation, dose, metabolism, animal species used, the company obtained from, sex, and fasting state (11–16). Further, clinical translation from rodents to primates is often lost due to discrepant findings that exist among preclinical models (17,18). Therefore, a more comprehensive understanding of the distribution of the canonical and extended ECS among preclinical animal models is necessary to increase scientific rigor and provide critical insight into the mechanisms by which phytocannabinoids elicit unexpected or seemingly contradictory findings across research groups.

To address this, we determined the relative expression of the 14 canonical and extended ECS genes (adgrf1, *adora2a, cnr1, cnr2, gpr18, gpr55, gpr119, faah, htr1a, naaa, ppara, pparg, trpv1,* and *trpv2)* in seven peripheral organs with metabolic and/or immune functions (colon, heart, kidney, liver, mesenteric lymph node [MLN], spleen, and visceral fat) in three translationally relevant preclinical animal models: C57BL/6 mice (*Mus musculus*), Sprague Dawley rats (*Rattus norvegicus*), and rhesus macaques (*Macaca mulatta*). Of note, while our present focus was on ECS relative gene distribution in the periphery, a subsequent publication will characterize distribution across sub-anatomic brain regions of these same animals.

## Materials and Methods

### Ethics statement

Animals and procedures in this study were approved by the Johns Hopkins University Animal Care and Use Committee. Animal handling and euthanasia were conducted as stated under the NIH Guide for the Care and Use of Laboratory Animals and USDA Animal Welfare Regulation. Rats and NHP included in this study were healthy uninfected animals serving as controls in other experiments (13).

#### 2.1 Animal use

##### 2.1.1 Mice

Five C57/BL6 mice (female [n=3] and male [n=2]) were included in this study. Mice were housed in ventilated racks with a 14/10-hours light/dark cycle, with water and standard chow diet (Teklad Diet 2018; IN, USA) *ad libitum*. Mice were kept in their cages for 13-weeks before they were sedated with isoflurane and euthanized. During necropsy, colon, heart, kidney, liver, spleen, and visceral fat tissue were collected, washed with 1X PBS to remove contaminating blood, and were flash frozen with liquid nitrogen and stored at −80°C until further use. No MLN was included in this study due to the complexity of identifying them due to their small size and dissecting both the brain and the periphery at the time of necropsy.

##### 2.1.2 Rats

Six female Sprague-Dawley rats (Charles River, MA, USA) were single-housed in wire-topped plastic cages in temperature and humidity-controlled facilities with a reverse light cycle (12 hours, lights off from 8:00am-8:00pm). Animals were provided corn-based chow (Teklad Diet 2018; IN, USA), and water *ad libitum*, except when actively participating as control subjects in behavioral procedures (13). Rats were 52-weeks old at the end of the study. Before necropsy, rats were sedated with isoflurane and euthanized by rapid decapitation. Upon necropsy, pancreas was the first organ to be collected and flash frozen. Afterwards, colon, heart, kidney liver, MLN, and spleen were collected, washed with fresh 1X PBS, flash frozen using dry ice and stored at −80°C until further use. No fat tissue was included in this study.

##### 2.1.3 Non-Human Primates

Four adult, male, pathogen-free Rhesus macaques (RM) (*Macaca mulatta*) were included in this study (animal identification numbers 560, 561, 562, and 563). Female macaques were not included in this study due to their importance in breeding for maintaining the colony. Macaques were pair-house to minimize any immunologic stress caused by being single-housed and they were fed standard monkey chow (Teklad Diet 2018; IN, USA) (19). Macaques were 7.89, 8.76, 8.59 and 7.95 years old at time of necropsy. During necropsy, animals were sedated using ketamine, and euthanized with an overdose of sodium pentobarbital, according to the American Veterinary Medical Association guidelines (2013). Phosphate buffered saline (1X) was used to perfuse organs and remove blood from organs, tissues, and MLN were taken to analyze the relative expression of the canonical, and the extended endocannabinoid receptors. No colon samples were available at the time of the study.

#### 2.2 RNA extraction & cDNA synthesis

RNA was extracted using RNeasy kit (Qiagen, MD, USA) following manufacturer’s instructions. Briefly, ∼200mg of each tissue were added to tubes containing Lysing Matrix D (MP Biomedicals, CA, USA). Tissue was homogenized using MP FastPrep®-24 (MP Biomedicals, CA, USA). Fat tissue was centrifuged after homogenization to remove the top layer of fat as instructed by the manufacturer. Afterwards, the aqueous phase was mixed with 70% ethanol (The Warner Graham Company, MD, USA) at a 1:1 ratio in a clean tube and loaded into the RNeasy columns. RNA-free DNase (Qiagen, CA, USA) was added to the column to digest any DNA present in the sample, as suggested by the manufacturer. RNA concentration and quality parameters were determined using Nanodrop (ThermoFisher Scientific, MA, USA). Extracted RNA was used to synthesize cDNA using iScript cDNA Synthesis kit (Bio-Rad, CA, USA) following manufacturer’s instructions.

#### 2.3 Real-Time Quantitative-Polymerase Chain Reaction

Relative genetic expression was determined using Real Time Quantitative Polymerase Chain Reaction (RT-qPCR) (CFX96™ Real-Time System, Bio-Rad, CA, USA) using commercially available TaqMan primers (**Tables 1-3**) and TaqMan™ Fast Universal PCR Master Mix (2X) no AMPERASE™ UNG (ThermoFisher Scientific, Catalog#4367846, MA, USA). Amplification was done in 40 cycles with the following conditions (Denaturing at 95°C for 20 seconds and annealing and extending at 60°C for 20 seconds). Cycle threshold values were normalized using Pan Eukaryotic 18S (ThermoFisher Scientific, MA, USA), transformed using the 2^-ΔCT^ method, and graphed to represent the relative genetic expression by sample, gene group and species.

**Table-1:**
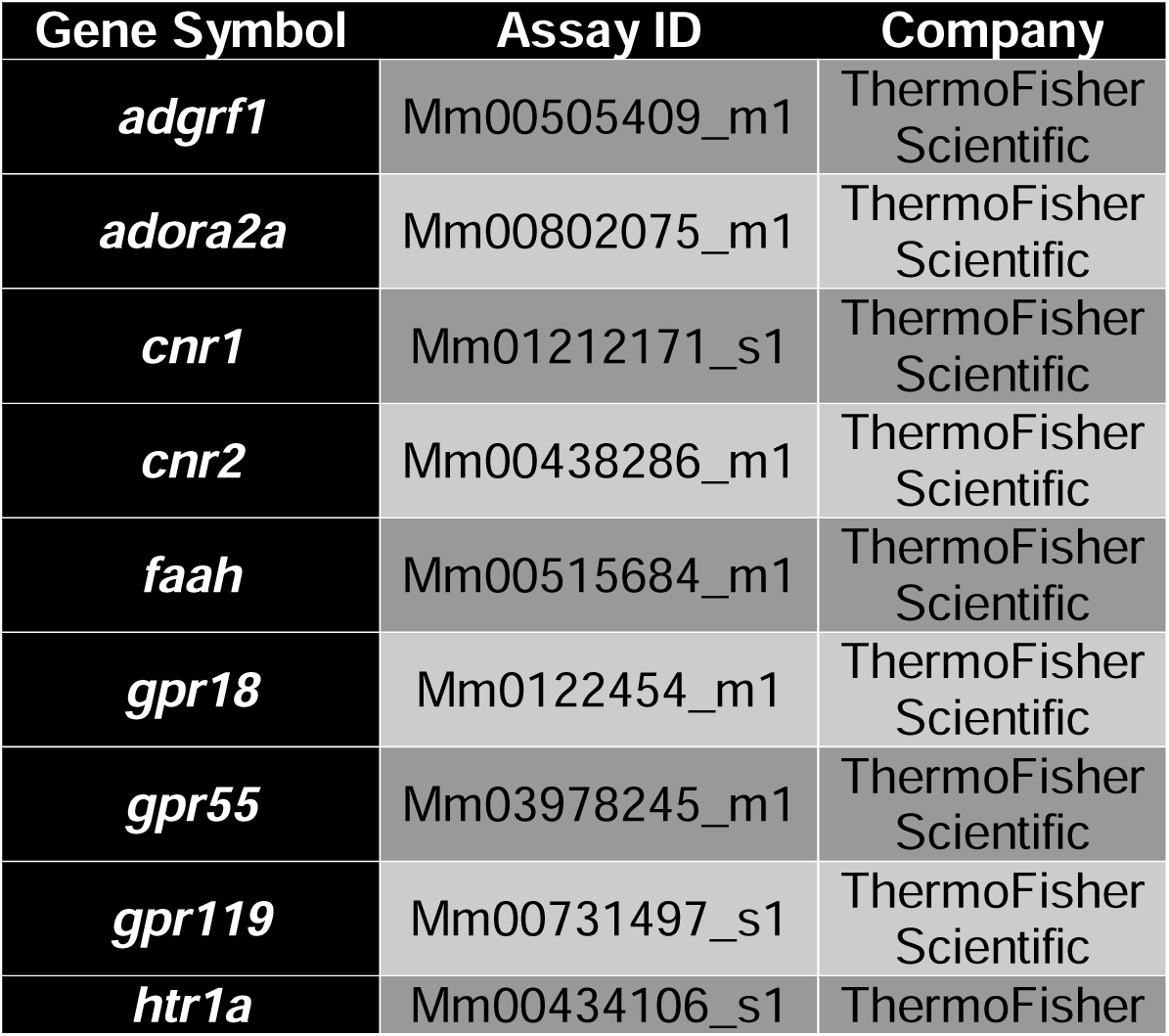
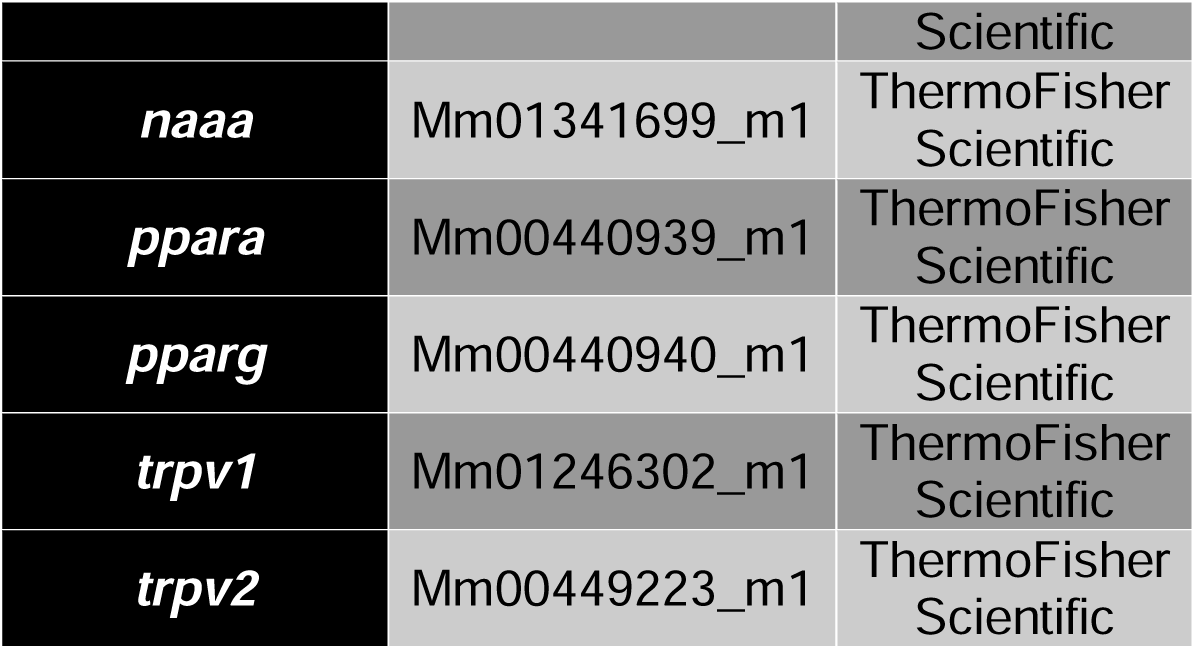
List of primers used to determine the relative expression of canonical and extended endocannabinoid receptors in mice (*Mus musculus*).

**Table-2:** List of primers used to determine the relative expression of canonical and extended endocannabinoid receptors in rats (*Ratus norvegicus*).

| **Gene Symbol** | **Assay ID** | **Company** |
| --- | --- | --- |
| ***adgrf1*** | Rn01511909_m1 | ThermoFisher Scientific |
| ***adora2a*** | Rn00583935_m1 | ThermoFisher Scientific |
| ***cnr1*** | Rn03037213_s1 | ThermoFisher Scientific |
| ***cnr2*** | Rn01637601_m1 | ThermoFisher Scientific |
| ***faah*** | Rn00577086_m1 | ThermoFisher Scientific |
| ***gpr18*** | Rn01493247_m1 | ThermoFisher Scientific |
| ***gpr55*** | Rn03037213_s1 | ThermoFisher Scientific |
| ***gpr119*** | Rn01648212_m1 | ThermoFisher Scientific |
| ***htr1a*** | Rn01637601_m1 | ThermoFisher Scientific |
| ***naaa*** | Rn01768319_m1 | ThermoFisher Scientific |
| ***ppara*** | Rn00566193_m1 | ThermoFisher Scientific |
| ***pparg*** | Rn00440945_m1 | ThermoFisher Scientific |
| ***trpv1*** | Rn00583117_m1 | ThermoFisher Scientific |
| ***trpv2*** | Rn00567974_m1 | ThermoFisher Scientific |

**Table-3:**
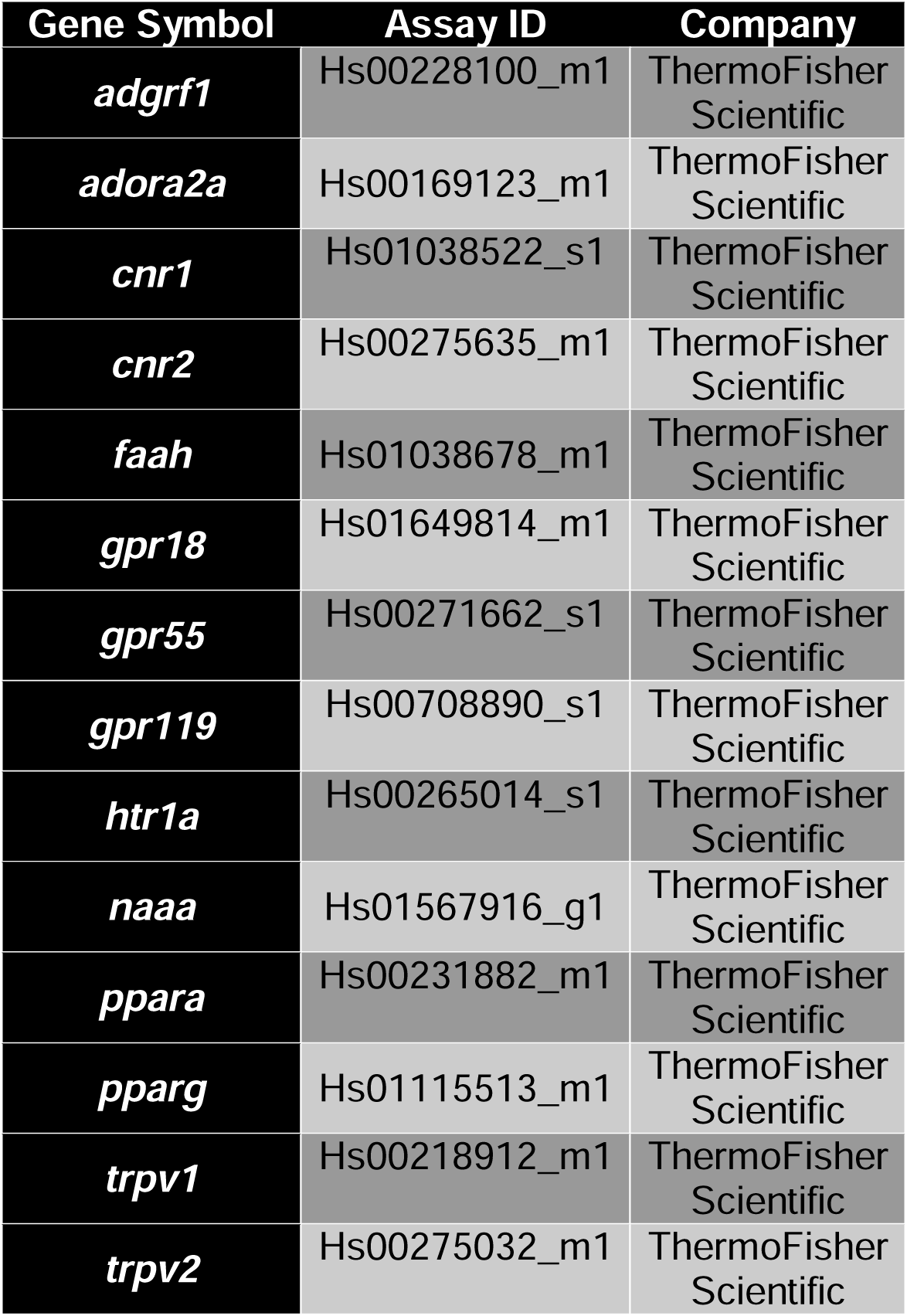
List of primers used to determine the relative expression of canonical and extended endocannabinoid receptors in Rhesus macaques (*Macaca mulatta*).

| Gene Symbol | Assay ID | Company |
| --- | --- | --- |
| *adgrf1* | Hs00228100_m1 | ThermoFisher Scientific |
| *adora2a* | Hs00169123_m1 | ThermoFisher Scientific |
| *cnr1* | Hs01038522_s1 | ThermoFisher Scientific |
| *cnr2* | Hs00275635_m1 | ThermoFisher Scientific |
| *faah* | Hs01038678_m1 | ThermoFisher Scientific |
| *gpr18* | Hs01649814_m1 | ThermoFisher Scientific |
| *gpr55* | Hs00271662_s1 | ThermoFisher Scientific |
| *gpr119* | Hs00708890_s1 | ThermoFisher Scientific |
| *htr1a* | Hs00265014_s1 | ThermoFisher Scientific |
| *naaa* | Hs01567916_g1 | ThermoFisher Scientific |
| *ppara* | Hs00231882_m1 | ThermoFisher Scientific |
| *pparg* | Hs01115513_m1 | ThermoFisher Scientific |
| *trpv1* | Hs00218912_m1 | ThermoFisher Scientific |
| *trpv2* | Hs00275032_m1 | ThermoFisher Scientific |

#### 2.4 Data Analysis and Statistics

Data were analyzed using PRISM software version 9.0 (GraphPad Software, Inc., San Diego, CA). Determination of relative gene expression was done in duplicates and represented in graphs plotting the mean±SEM. Our limit of detection (LoD) was calculated using an average of each species probing for Pan Eukaryotic 18S (ThermoFisher Scientific, MA USA) with a cycle threshold of 35. Please note that each sample was subtracted each own 18S value and hence can appear below the LoD but its expression was detected in a Ct value below 35. Samples that did not amplify were given an arbitrary value of 39.99. Variance between the relative expression of genes between organs and by species was determined using one-way ANOVA. Post hoc analysis was done to determine the difference between the expression of these genes when there was statistical significance determined by one-way ANOVA. (*p≦0.05, **p≦0.01, ***p≦0.001 & ****p≦0.0001)

## Results

### Both Canonical ECS Receptors Are Present In The Spleen Of Mice, Rats and NHP

*Cnr1* is primarily expressed and studied in the context of the brain (20,21). Here we show that *cnr1* mRNA was detected in both metabolic and secondary immune organs in mice, including the colon, kidney, spleen, and visceral fat (**Figure-1A)** Interestingly, significantly more *cnr1* was present in the visceral fat as compared to all other evaluated organs (p=0.0277 vs. colon, 0.0250 vs. heart, 0.0250 vs. kidney, 0.0250 vs. liver and p=0.302 vs. spleen). In contrast to mice, *cnr1* was more restricted in rats where the highest levels occurred in kidney (**Figure-1B**). Indeed, *cnr1* was significantly higher in kidney as compared to heart (p-value=0.0257), liver (p-value=0.0209), MLN (p-value=0.0271), and spleen (p-value=0.0304). While *cnr1* was present in colon, MLN, and spleen, it did not occur in all rats with only 4/6, 1/6, and 1/6 rats having detectable expression, respectively. *Cnr1* was least abundant in NHP, where it was limited to the spleen and the visceral fat (**Figure 1C**). *Cnr1* was not detectable in the liver or heart in any of the three models evaluated in this study.

**Figure-1:**
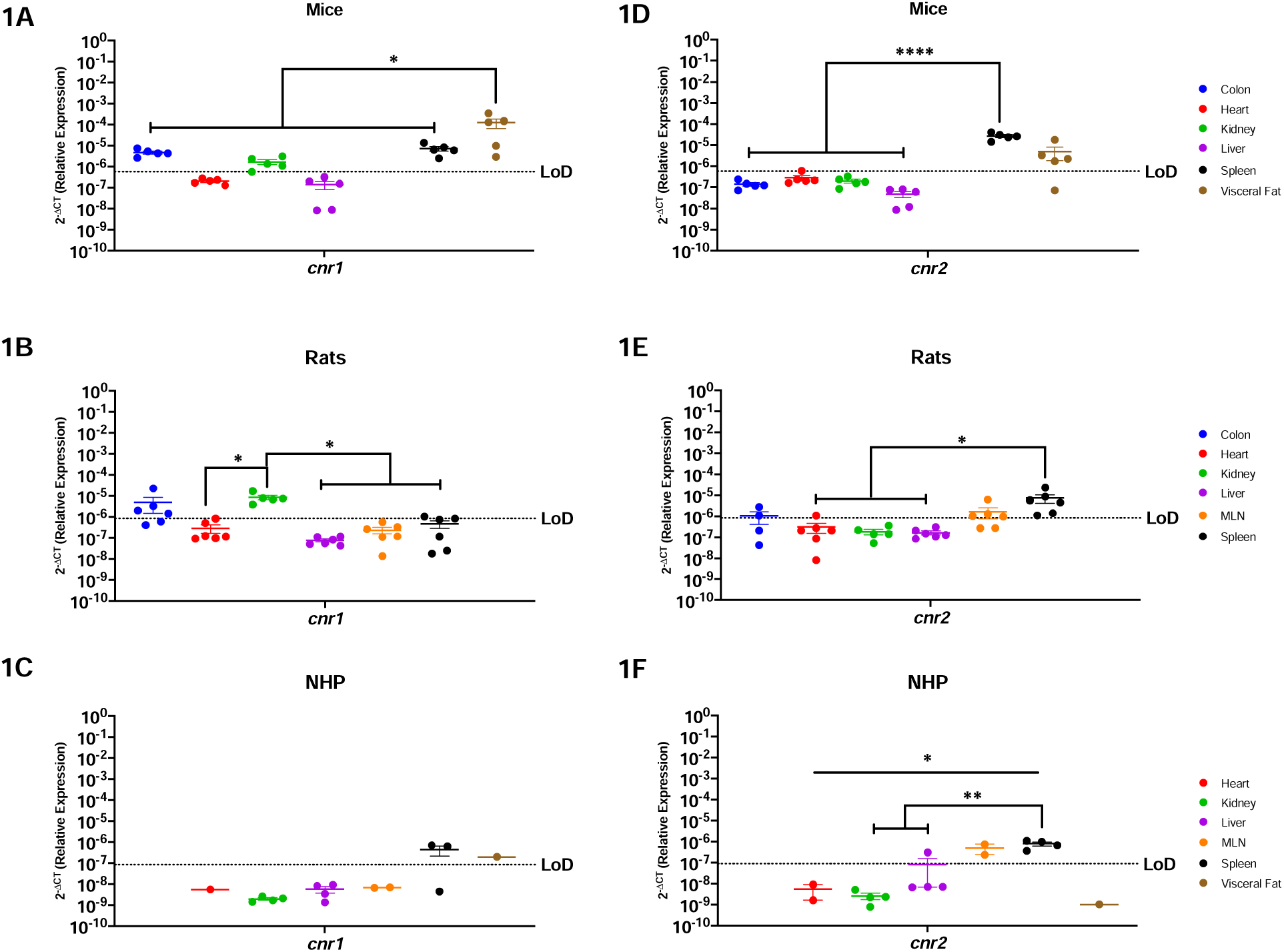
Both Canonical ECS Receptors Are Present In The Spleen Of Mice, Rats and NHP. Relative expression of *cnr1* and *cnr2* was determined using qPCR from the colon, heart, kidney, liver, MLN, spleen, and visceral fat from mice, rats and NHP. **A)** C*nr1* mRNA in mice was detected in the colon, kidney (4/5 mice), spleen, and visceral fat, having the highest levels in the latter. **B)** C*nr1* was detected in partial samples of the rat model colon (4/6 rats), heart (1/6 rats), kidney, and spleen (1/6 rats), having statistically higher levels of expression in the kidney when compared to the heart, liver, MLN, and spleen. **C)** In NHP, *cnr1* mRNA was detected in the spleen (2/3 rhesus) and visceral fat at comparable levels. **D)** *Cnr2* was detected only in the spleen and visceral fat (4/5 mice), at comparable levels. The spleen had statistically significantly higher levels as compared to the colon, heart, kidney, and liver. **E)** *Cnr2* was detected in the colon (2/4 rats), heart (1/6 rats), MLN (4/6 rats) and spleen of rats. **F)** In NHP, *cnr2* mRNA was detected in the liver (1/4 rhesus), MLN and spleen. Detection levels were significantly higher in the spleen when compared to the heart, kidney, and liver, but with significant differences when compared to the liver (1/4 rhesus), kidney, and heart. Data are represented as the mean ± SEM. (*p-value >0.05, **p-value>0.001, ***p-value>0.0001, ****p-value>0.00001).

*Cnr2* is primarily present in the periphery with expression in the brain occurring in the context of disease (20–22). Our findings were consistent with this, where *cnr2* mRNA was detected in the spleen and in the visceral fat (4/5 mice) of mice at similar levels. Significant differences were found among the spleen and the colon (p=<0.0001), heart (p=<0.0001), kidney (p=<0.0001), and liver (p=<0.0001). **(Figure-1D)**. In rats, we detected *cnr2* mRNA partially in the colon (2/4 rats), the heart (1/6 rats), MLN (4/6 rats), and the spleen (6/6 rats) **(Figure-1E)**. Significant differences were found among the spleen and the heart (p=0.0237), kidney (p=0.0299), and liver (p=0.0201). NHP had restricted *cnr2* mRNA, with robust levels in the MLN and spleen **(Figure-1F)**. *Cnr2* was partially detected in the liver (1/4 rhesus) in the NHP model. Notably, *Cnr2* in the spleen was significantly more highly expressed in NHP when compared to the heart (p=0.0182), kidney (p=0.0043) and liver (p=0.0092).

### Peroxisome Proliferator Activated Receptors, *Ppara* and *Pparg,* Are Generally Well Conserved In All Organs Of Mice, Rats and NHP

Peroxisome proliferator activated receptors mediate several vital functions, and hence are known to be expressed almost ubiquitously (23–25). Our findings corroborated this, where we determined that *ppara* mRNA was present in every evaluated organ in mice, with the notable exception of the spleen (**Figure 2A**). These genes were most abundantly expressed in the heart (p=0.0049), kidney (p=0.0484), and liver (p=0.0049). Similar trends occurred in rats, where *ppara* mRNA was detected in all organs available, having significantly higher expression in the liver vs. secondary immune organs (p=0.0261 for MLN, and p=0.0233 for spleen) and the colon (p=0.0445) **(Figure-2B)**. The NHP model also showed ubiquitous *ppara*, being detected in all evaluated organs **(Figure-2C)**.

**Figure-2:**
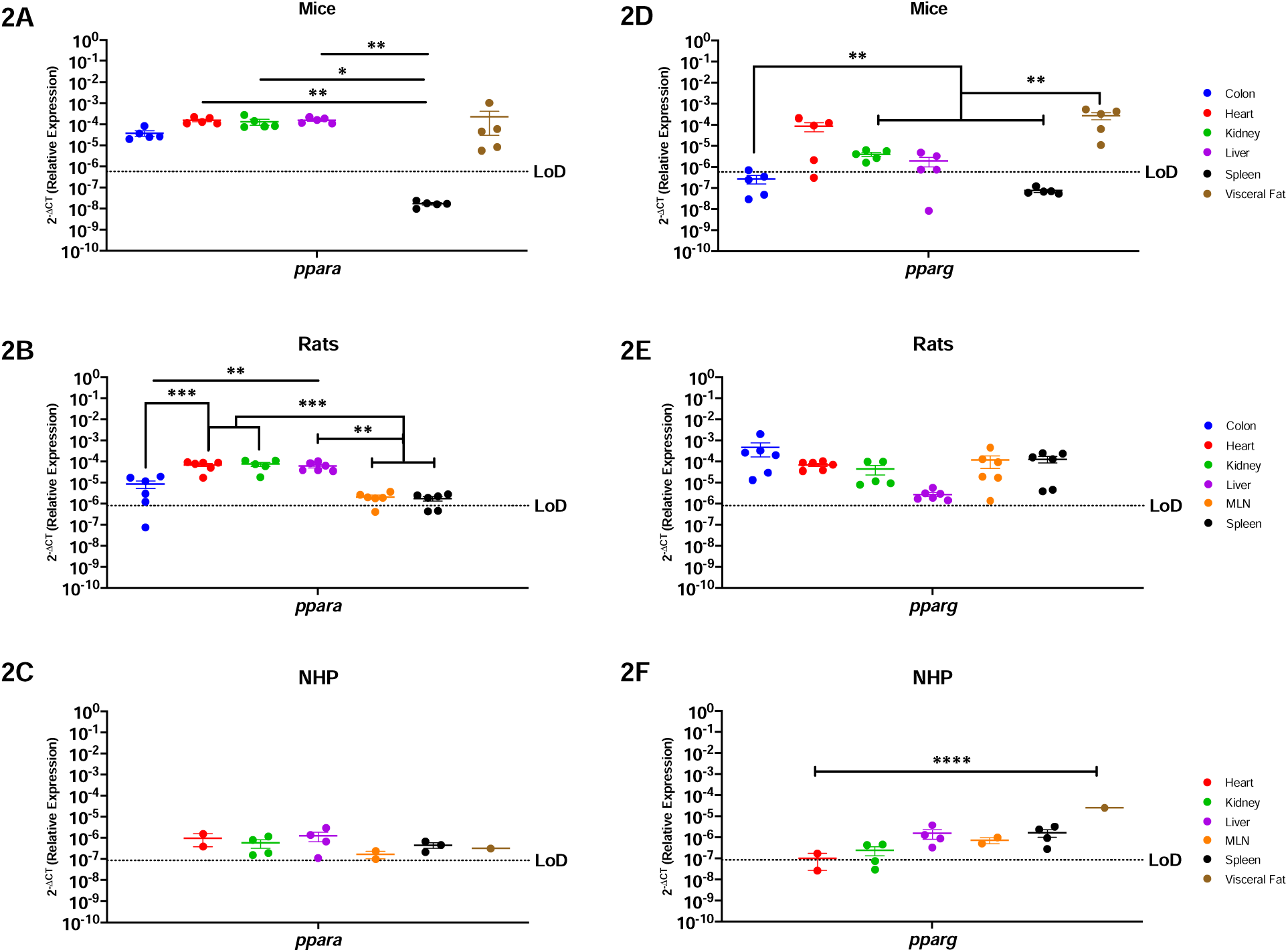
Peroxisome Proliferator Activated Receptors, *Ppara* and *Pparg,* Are Generally Well Conserved In All Organs Of Mice, Rats and NHP. Relative expression of *ppara* and *pparg* was determined using qPCR from the colon, heart, kidney, liver, MLN, spleen, and visceral fat from mice, rats, and NHP. **A)** P*para* mRNA was detected in all the organs tested in mice, except for the spleen. It had significantly higher levels in the kidney when compared with the colon and the spleen. **B)** In rats, *ppara* mRNA was detected in the colon (4/6 rats), heart, liver, kidney, MLN (5/6 rats), and spleen (4/6 rats). P*para* in the rat model was significantly higher in the heart, kidney, and liver, when compared to the colon, MLN, and spleen. **C)** *Ppara* was detected in all organs tested in the periphery of the NHP model at comparable levels for all the evaluated organs. **D)** Mouse *pparg* was similar to *ppara*, being detected in the colon (3/5 mice), heart (4/5 mice), kidney, liver (4/5 mice) and the visceral fat. Detection of this gene was higher in visceral fat when compared to the colon, kidney, liver, and spleen. **E)** *Pparg* was detected in all the organs available for the rat model with no statistical significance among any organ. **F)** P*parg* in the NHP model was also detected in all available organs for the NHP model, having only partial detection in the heart (1/2 rhesus) and the kidney (3/4 rhesus). Interestingly, *pparg* was significantly higher in the visceral fat when compared with all other tissues. It is worth mentioning that neither of these genes were detected in the spleen of mice, contrary to the other animal models. Data are graphed as the geometric mean ± SEM (*p-value >0.05, **p-value>0.001, ***p-value>0.0001, ****p-value>0.00001).

*Pparg* mRNA followed a similar trend as *ppara* in mice, where it was detected in all organs except for the spleen **(Figure-2D)**. Interestingly, *pparg* was most highly expressed in visceral fat and heart in mice. Significant differences were found among the visceral fat and colon (p=0.0027), kidney (p=0.0031), liver (p=0.0028), and spleen (p=0.0026). *Pparg* mRNA was also expressed across all organs in rats, where no significant differences occurred across the body **(Figure-2E)**. In contrast, NHP had nuanced *pparg* expression where it was highly present in visceral fat, when compared to the heart (p=0.0001), kidney (p=0.0001) liver (p=0.0001), MLN (p=0.0001), and spleen (p=0.0001) **(Figure-2F)**. In sum, *ppara* and *pparg* were similarly present in all animal models, with high expression detected in all organs, with the notable exception of the spleen of mice.

### Endocannabinoid-like GPRs Are Preferentially Expressed In Lymphoid Organs and The Visceral Fat

GPRs are mostly considered to be orphan receptors until identification of their specific ligand. Some GPRs (i.e., *gpr18*, *gpr55*, and *gpr119*) are known to interact with cannabinoids and are considered to be endocannabinoid-like GPRs (26–31). Mice had relatively limited *gpr18* mRNA, which was present only in the spleen and visceral fat **(Figure-3A)**. Surprisingly, rats had a very different profile where *gpr18* was detected among all organs, except for liver **(Figure-3B)**. Notably, *gpr18* was most highly expressed in the secondary lymphoid organs in rats, being statistically different from the colon (p=0.0120), heart (p=0.0137), kidney (p=0.0255), and liver (p=0.0092). *Gpr18* was only partially detected in the spleen (3/4 rhesus) in the NHP model **(Figure-3C)**.

**Figure-3:**
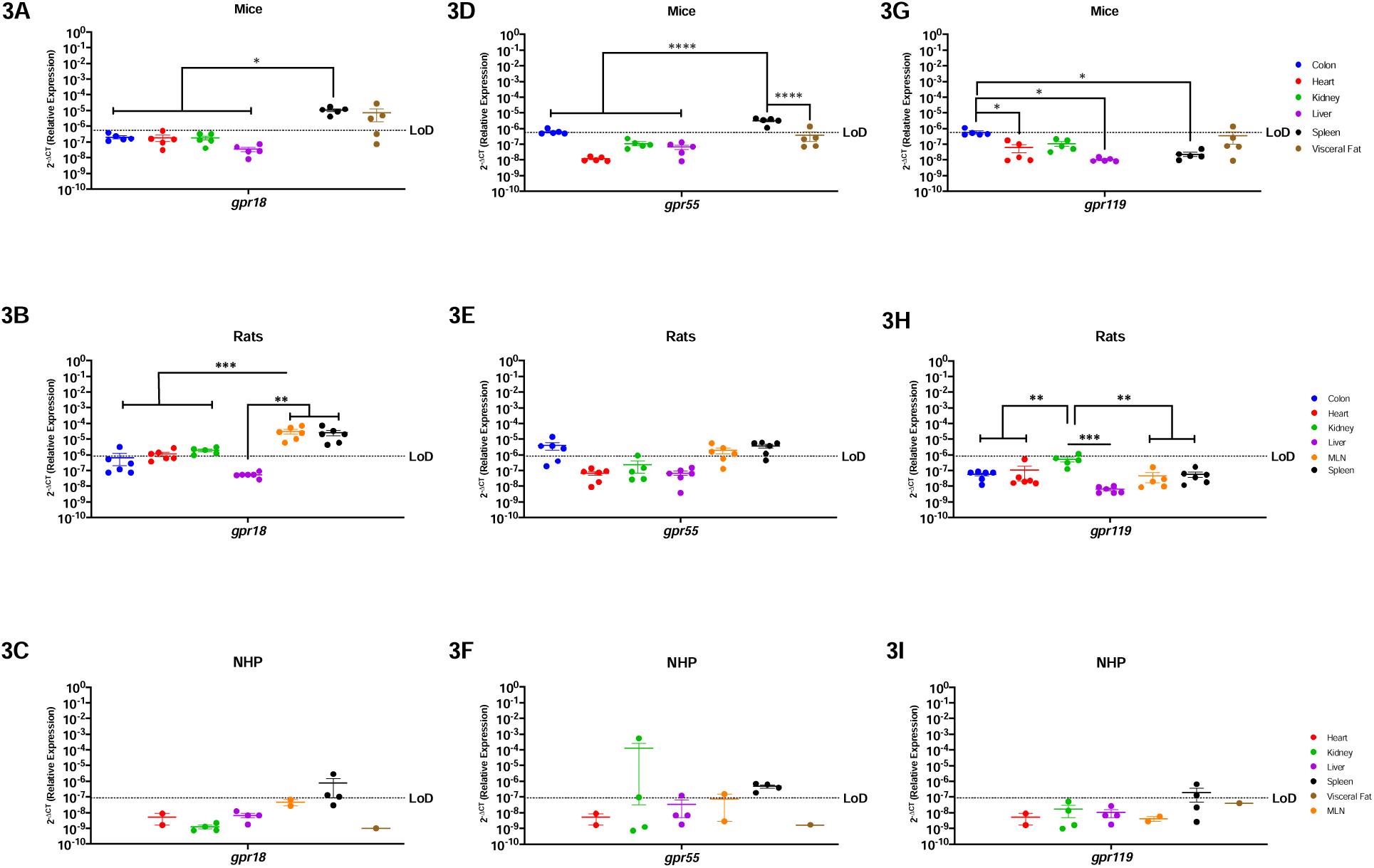
Endocannabinoid-like GPRs Are Preferentially Expressed In Lymphoid Organs and The Visceral Fat. Relative expression of *gpr18, gpr55,* and *gpr119* was determined using qPCR from the colon, heart, kidney, liver, MLN, spleen, and visceral fat from mice, rats and NHP. **A)** *Gpr18* was detected in the spleen and visceral fat with significant difference between the spleen and the rest of the organs. **B)** In rats, *gpr18* was highest in the secondary immune organs (MLN and spleen). **C)** *Gpr18* was only detected partially in the spleen (3/4 rhesus) of the NHP model. **D)** *Gpr55* was detected in the colon (1/5 mice), spleen and visceral fat (1/5 mice), having statistical significance among the spleen and all other tissues. **E)** *Gpr55* expression was detected in the colon (4/6 animals), MLN (4/6 rats) and the spleen (5/6 rats). **F)** In the NHP model, *gpr55* was detected in the kidney (1/4 rhesus), liver (1/4 rhesus), MLN (1/2 rhesus), and spleen (2/4 rhesus). **G)** *Gpr119* was detected partially in the colon (1/5 mice) and the visceral fat (1/5 mice), having statistical significance between the colon and the heart, liver, and spleen. **H)** *Gpr119* was only detected in the kidney (1/6 rats) with significant differences with all the other organs included in this study. **I)** *Gpr119* was partially detected in the spleen (2/4 rhesus) of the NHP model. Data represents the geometric mean ± SEM (*p-value >0.05, **p-value>0.001, ***p-value>0.0001, ****p-value>0.00001).

G*pr55* was similar to *gpr18* in mice, being highly expressed in the spleen and having significantly higher levels when compared to the colon (p=0.0414), heart (p=0.0278), kidney (p=0.0272), liver (p=0.0275), and visceral fat (p=0.0027) **(Figure-3D)**. *Gpr55* also primarily followed a similar expression pattern as *gpr18* in rats; however, they did not express *gpr55* in the heart or liver **(Figure-3E)**. Interestingly, NHP *gpr55* was partially detected in the kidney (2/4 animals), liver (1/4 animals), and MLN (1/2 animals), showing more expression than the other two endocannabinoid-like GPCRs, and a different pattern than the rodent *gpr18* and *gpr55* **(Figure-3F)**.

*Gpr119* generally followed a similar pattern to *gpr18* where it was minimally detected in all animal models. *Gpr119* was only present in the colon and visceral fat of mice **(Figure-3G)**, was undetectable in the periphery of rats **(Figure-3H)**, and only partially detected in the spleen of NHP (2/4 animals) **(Figure-3I)**.

### TRPV1 and TRPV2 Nociception Channels Have Limited Translational Applicability to Humans

Nociception channels are widely studied in their response to painful stimuli (32,33). *Trpv1* mRNA was detected in the colon, kidney, and visceral fat of mice, having significant differences when comparing the visceral fat to the colon (p=0.0076), heart (p=0.0050), kidney (p=0.0139), liver (p=0.0044) and spleen (p=0.0036) **(Figure-4A)**. In contrast, rats had a wider expression of this gene, which was detected in all evaluated organs **(Figure-4B)**. *Trpv1* was highest in the rat kidney and statistically increased as compared to the heart (p=0.0038), liver (p=0.0039), MLN (p=0.0039) and spleen (p=0.0041). NHP were comparable to mice and only had detectable *trpv1* in the kidney and spleen **(Figure-4C)**.

**Figure-4:**
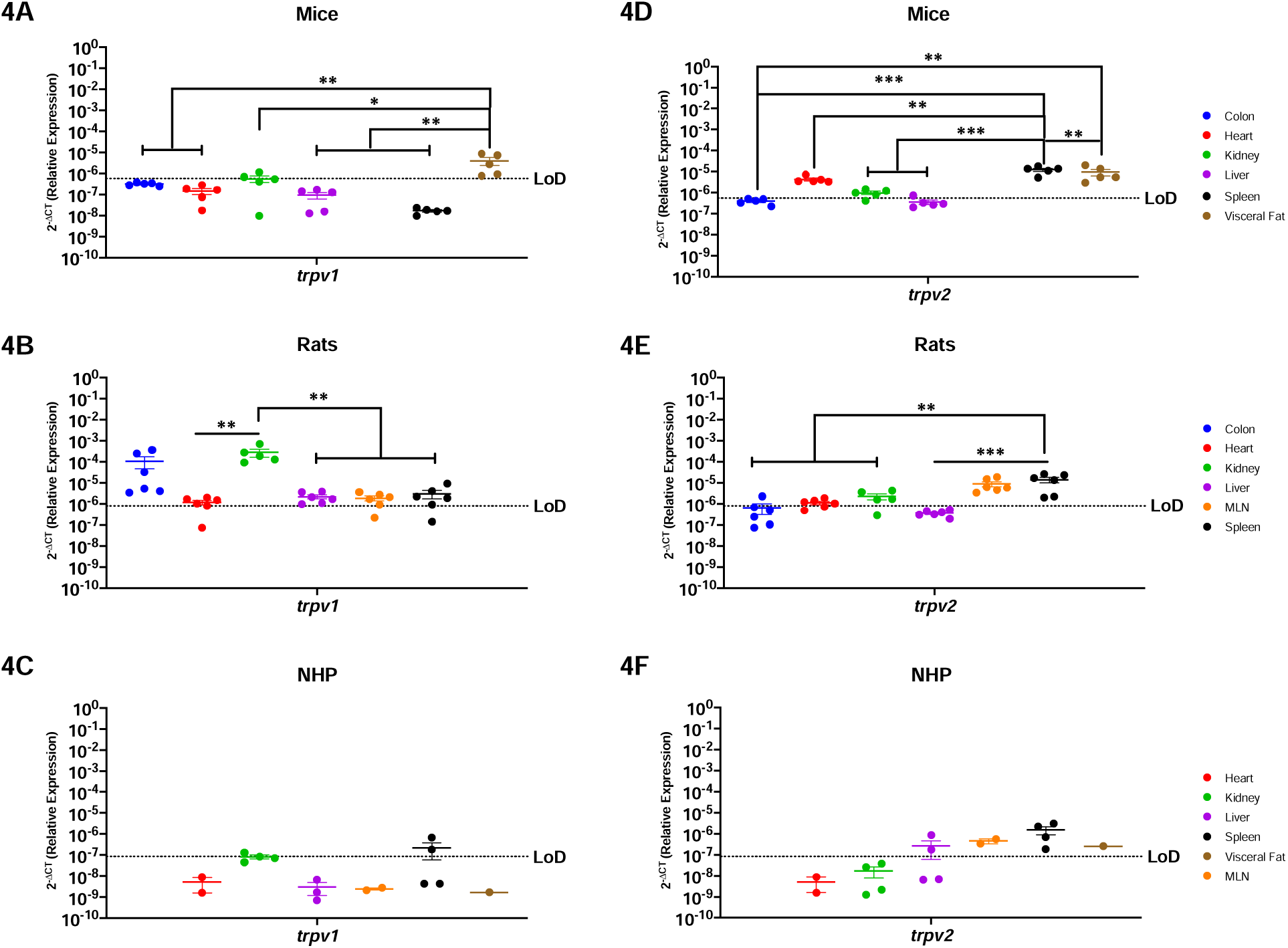
Peripheral TRPV1 and TRPV2 Nociception Channels Have Limited Translational Applicability to Humans. Relative expression of *trpv1* and *trpv2* was determined using qPCR from the colon, heart, kidney, liver, MLN, spleen, and visceral fat from mice, rats and NHP. **A)** *Trpv1* was detected in the colon, partially in the kidney (3/5 mice), and the visceral fat. **B)** *Trpv1* was detected in the colon, heart (5/6 animals), kidney, liver, MLN (5/6 rats) and spleen (5/6 rats). Significant differences were found between the kidney and the other organs, except for the colon which showed comparable expression levels. **C)** *Trpv1* was detected in the kidney (1/4 rhesus) and the spleen (2/4 rhesus). **D)** *Trpv2* mRNA was detected in the heart, kidney (4/5 mice), liver (1/6 mice), spleen, and visceral fat, having higher significant levels in the last two. **E)** *Trpv2* was detected in the colon (1/6 rats), heart (4/6 rats), kidney (4/5 rats), MLN, and spleen. T*rpv2* was similar between immune organs, and they are both significantly different when compared with the metabolic organs. **F)** *Trpv2* showed broader detection, being detected in the liver (2/4 rhesus), MLN, spleen, and visceral fat. Data are graphed as the geometric mean ± SEM (*p-value >0.05, **p-value>0.001, ***p-value>0.0001, ****p-value>0.00001).

*Trpv2* mRNA was widely present in mouse with expression in the heart, kidney, liver, spleen, and visceral fat **(Figure-4D)**. Even still, *trpv2* was not expressed equally but instead was significantly higher in the spleen and the visceral fat as compared to colon (p=0.0002), heart (p=0.0117), kidney (p=0.0003), and liver (p=0.002). *Trpv2* was detected throughout all the organs in the rat, except for the liver **(Figure-4E)**. Of the organs in which it was expressed, *trpv2* was most highly present in the rat spleen, with comparable levels to MLN, but reaching statistical significance with the colon (p=0.0010), heart (p=0.0016), and kidney (p=0.0071). Interestingly, the NHP model was the only preclinical model where *trpv2* was present in liver **(Figure-4F)**. In contrast, *trpv2* was detectable in the NHP MLN, spleen, and visceral fat, and was comparably expressed across organs without any significant differences between them.

### Endocannabinoid Metabolic Enzymes Are Ubiquitous In Rodents, But More Restricted In NHP

*Faah* and *naaa* are ubiquitous enzymes involved in endocannabinoid degradation (34–38). In agreement with this, *faah* was present in all organs analyzed in mice, having more abundance in the kidney and the liver as compared to the colon (p=0.0067 & p>0.0001, respectively), heart (p=0.0013 & p>0.0001, respectively), visceral fat (p=0.0116 & p>0.0001, respectively), and the spleen, but only when compared to the liver (p=0.0021) **(Figure-5A).** Similarly, *faah* was present in all rat organs, having increased expression in the colon and decreased expression in the heart (p=0.0264), following a similar pattern as the mice **(Figure-5B)**. Interestingly, NHP had a different *faah* expression pattern than the rodents. While widely detected in the kidney, liver (3/4 rhesus), spleen (3/4 rhesus) and visceral fat, *faah* was not detected in the NHP heart nor MLN **(Figure-5C)**.

**Figure-5:**
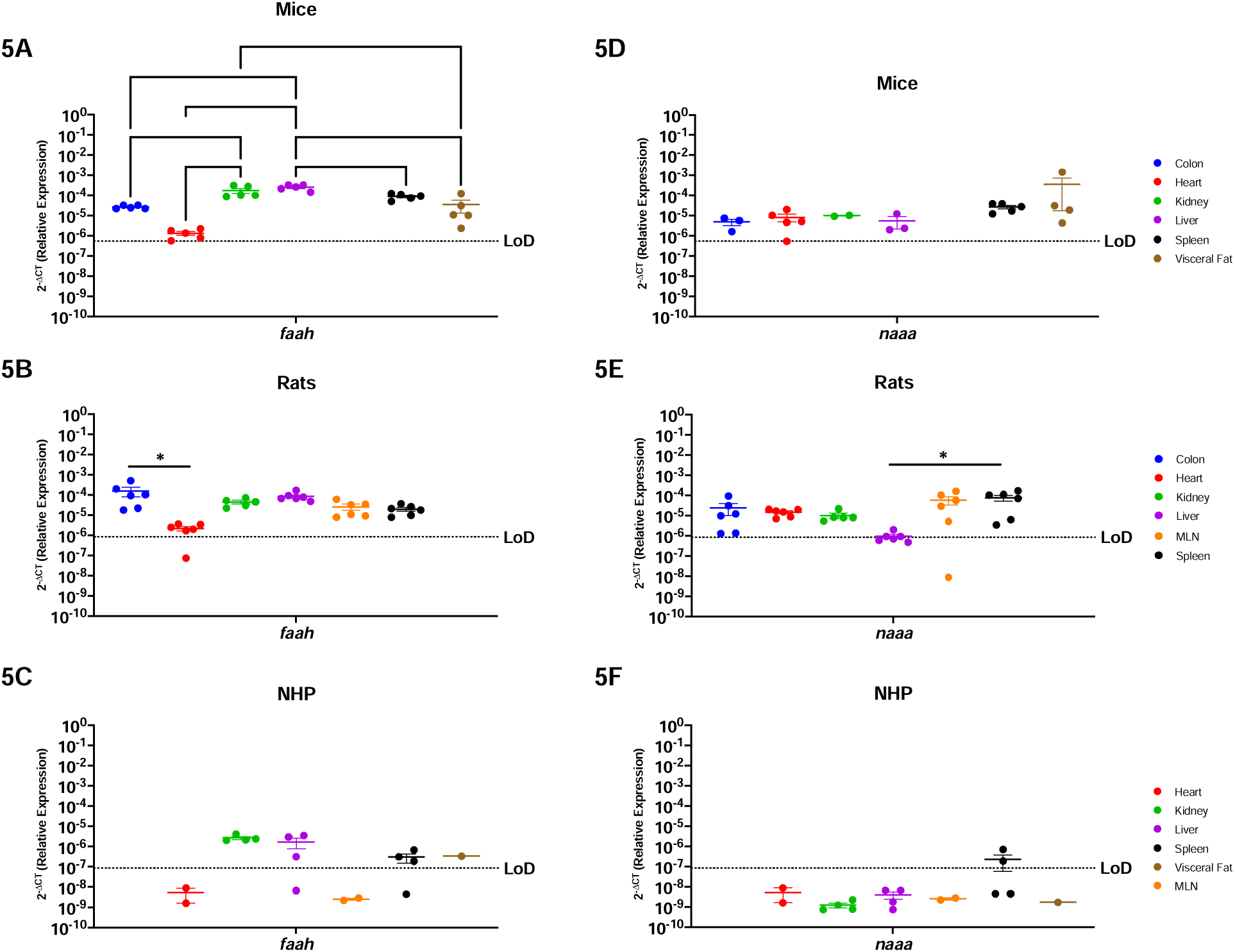
Endocannabinoid Metabolic Enzymes Are Ubiquitous In Rodents, But More Restricted In The NHP Model. Relative expression of *faah* and *naaa* was determined using qPCR from the colon, heart, kidney, liver, MLN, spleen, and visceral fat from mice, rats and NHP. **A)** F*aah* mRNA was detected in all organs tested, having differential expression between them. The heart of mice had the lowest levels for this gene. **B)** *Faah* mRNA was detected in all organs included in this study. The colon of rats showed higher levels of *faah*, particularly significant when compared to expression in the heart. **C)** Levels of *faah* in the NHP model were detected in the kidney, liver (3/4 animals), spleen (3/4 rhesus), and visceral fat. **D)** N*aaa* mRNA was detected in all organs with similar levels. **E)** Expression of *naaa* mRNA in rats was detected in all organs and was least abundant in liver. **F)** *Naaa* mRNA was detected only partially in the spleen (2/4 rhesus). Data are graphed as the geometric mean ± SEM (*p-value >0.05, **p-value>0.001, ***p-value>0.0001, ****p-value>0.00001).

*Naaa* showed similar trends as *faah* where rodents had ubiquitous expression while NHP had more restricted expression. *Naaa* was detected in all organs tested in mice, to comparable levels **(Figure-5D)**. In rats, it was also detected in all organs, but with significantly lower expression in the liver when compared to the spleen (p=0.0289) **(Figure-5E)**. *Naaa* was only expressed in the NHP spleen (2/4 rhesus).

### *Htr1a*, *Adora2a* and *Adgrf1* are Poorly Conserved Among Mice, Rats, and NHP

*Htr1a* is a serotonin receptor primarily studied in the brain, while *adora2* and *adgrf1* are more widely present and implicated in inflammation, cardiovascular diseases, and cancer (39–45). In general, *htr1a* mRNA was minimally expressed in the periphery of all three preclinical models. *Htr1a* was only detected in the visceral fat (1/5 mice), colon (1/5 rats), and liver (2/3 rhesus) and spleen (2/4 rhesus) **(Figure-6A-C)**. *Adora2a* is present to a greater extent in the periphery and was detected in the heart, kidney, spleen, and visceral fat of mice **(Figure-6D)**. Rats also widely expressed *adora2a* which was found in all examined organs but was lowly expressed in the colon (p=0.0014) and the heart (p=0.0384) **(Figure-6E)**. NHP also expressed *adora2a*, however it was present only in the spleen and MLN secondary immune organs **(Figure-6F)**. *Adgrf1* was detected in the colon, kidney, liver, and visceral fat from mice, though it was mostly highly expressed in the liver **(Figure-6G)**. Rats had widespread *adgrf1* across all organs in rats, with preferential expression in the kidney when compared with the colon (p=0.0286), heart (p=0.0273), liver (p=0.0396), and the spleen (p=0.0472) **(Figure-6H)**. *Adgrf1* had the most limited expression in NHP where it was detected only in the kidney (1/4 rhesus) **(Figure-6I)**.

**Figure-6:**
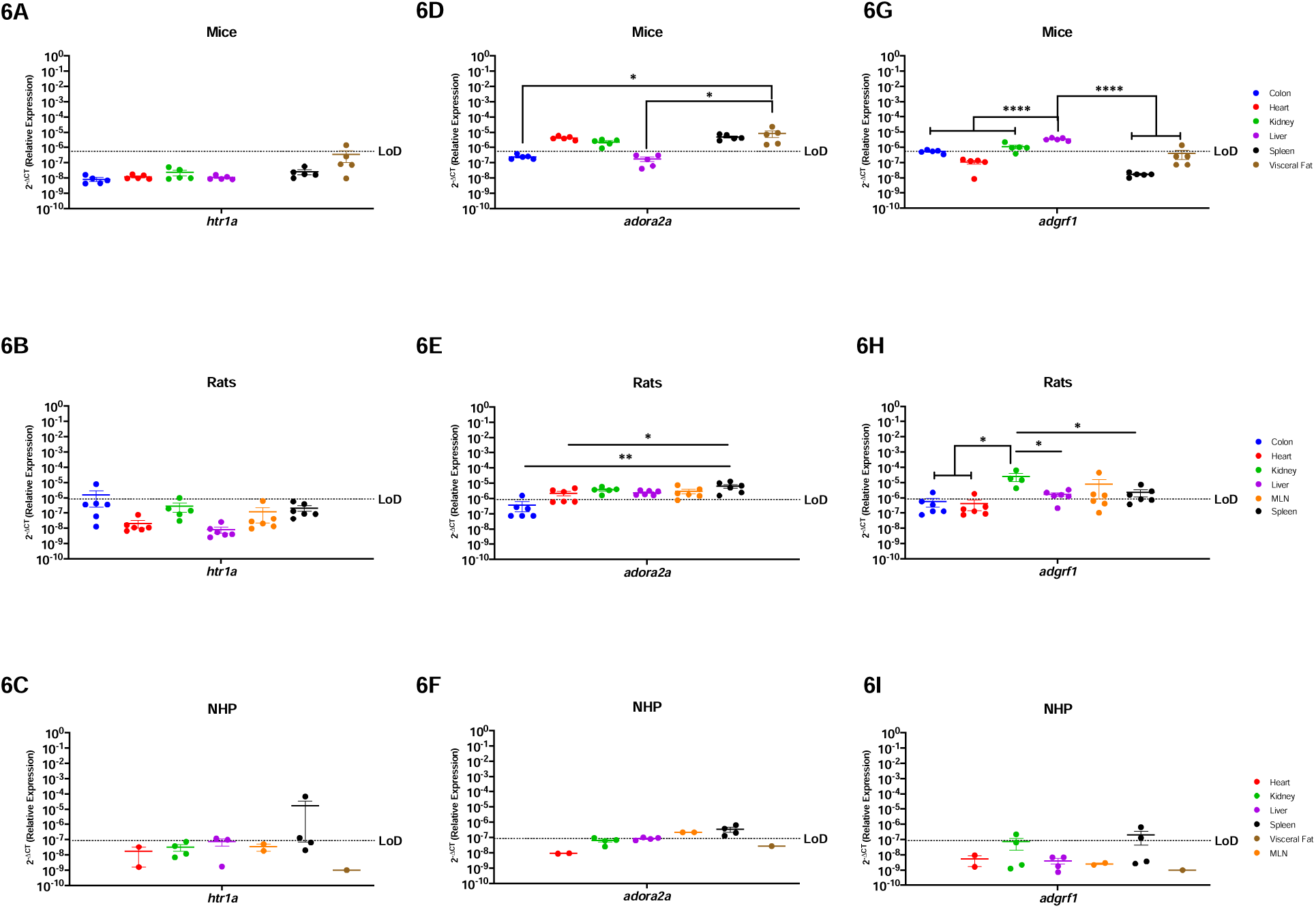
Htr1a, *Adora2a* and *Adgrf1* are Poorly Conserved Among Mice, Rats, and NHP. Relative expression of *5-htr1a*, *adora2a* and *adgrf1* was determined using qPCR from the colon, heart, kidney, liver, MLN, spleen, and visceral fat from mice, rats and NHP. **A)** *Htr1a* was only detected partially in the in the visceral fat of mice (1/6 mice). **B)** In rats, *htr1a* was only detected in one sample of the colon. **C)** H*tr1a* in the NHP model was detected in the spleen (2/4 rhesus). **D)** In mice, *Adora2a* was detected in the heart, kidney, spleen, and visceral fat. The visceral fat levels were significantly higher as compared to colon and liver. **E)** *Adora2a* was detected in all organs screened, with significantly higher levels in the spleen when compared to other organs. **F)** A*dora2a* in NHP was detected in the MLN and spleen. **G)** *Adgrf1* was detected in the colon, kidney, liver, and visceral fat (1/6 mice). This gene was higher in the liver when compared to all the other organs. **H)** *Adgrf1* was detected in the kidney with significant higher levels when compared to the colon, liver, and spleen. This gene was also detected in the heart (1/6 rats) and MLN (3/5 rats) **I)** *Adgrf1* was detected partially in the kidney (1/4 rhesus) and the spleen (2/4 rhesus). Data are graphed as the geometric mean ± SEM (*p-value >0.05, **p-value>0.001, ***p-value>0.0001, ****p-value>0.00001).

## Discussion

We report a comparison of the relative expression of 14 genes from the canonical and extended ECS in seven peripheral organs from three animal species and strains widely used in research, including in cannabis and cannabinoid research: C57BL/6 mice, Sprague-Dawley rats, and Rhesus macaque NHP. We identified key differences in the relative expression patterns of these evolutionary conserved, polyfunctional receptors, and found that these preclinical model systems were more dissimilar than has been previously appreciated. Indeed, there were only five receptors (CB2, GPR18, GPR55, TRPV2, and FAAH) that had identical expression patterns in all three preclinical animal models. Of note, these five receptors were consistently expressed in the spleen for all species evaluated, suggesting the importance of endocannabinoid function in this secondary lymphoid organ and potential for therapeutic intervention. This indicates that while the ECS is highly conserved, each of the three animal species included has a differing pattern for receptor composition in their peripheral organs. This is incredibly important and has profound implications for translation to humans and for comparison across research groups. The impact of route of administration, diet, formulation, dose, fasting vs. fed state, biological sex, and metabolite distribution and bioavailability (11) is already implicated in contributing to discrepant findings in the cannabinoid field. We propose that the unique receptor composition patterns of the preclinical model must also be considered to enhance scientific rigor and reproducibility. Indeed, as multiple canonical and extended ECS receptors are simultaneously present within a tissue, the potential for off-target and polypharmacy effects (14,16,46–51,51–67) is staggering as each receptor has its own unique function and signaling processes. Therefore, it is important to understand the nuances of endocannabinoid receptor tissue localization in the most common preclinical animal models.

Our findings also bring attention to the importance of additional receptors that have been understudied thus far. While the canonical ECS receptors, CB1 and CB2, have been most widely studied, our work demonstrates that the extended ECS receptor distribution represents an additional level of complexity that must be considered when performing cannabinoid studies. Indeed, these receptors and/or metabolic enzymes are simultaneously present in peripheral tissues in tandem with CB1 and/or CB2. and are also capable of mediating physiologic effects upon interacting with endo- and phytocannabinoids. These interactions should not be ignored as they result in a complex network of physiological pathways having diverse effects in biologic systems in the chosen preclinical model. Our results are summarized in **Tables 4-6**.

**Table-4:** Summary of the findings of the expression of the canonical and extended ECS in mice (*Mus musculus*).

| Mice | Colon | Heart | Kidney | Liver | MLN | Spleen | Visceral Fat |
| --- | --- | --- | --- | --- | --- | --- | --- |
| *cnr1* | + | - | + | - |  | + | + |
| *cnr2* | - | - | - | - |  | + | + |
| *ppara* | + | + | + | + |  | - | + |
| *pparg* | + | + | + | + |  | - | + |
| *gpr18* | + | - | - | - |  | + | + |
| *gpr55* | + | - | - | - |  | + | + |
| *gpr119* | + | - | - | - |  | - | + |
| *trpv1* | + | - | + | - |  | - | + |
| *trpv2* | - | + | + | + |  | + | + |
| *faah* | + | + | + | + |  | + | + |
| *naaa* | + | + | + | + |  | + | + |
| *htr1a* | - | - | - | - |  | - | + |
| *adora2A* | - | + | + | - |  | + | + |
| ***adgrf1*** | + | - | + | + |  | - | + |

**Table-5:** Summary of the findings of the expression of the canonical and extended ECS in rats (*Ratus norvegicus*).

| Rat | Colon | Heart | Kidney | Liver | MLN | Spleen | Visceral Fat |
| --- | --- | --- | --- | --- | --- | --- | --- |
| ***cnr1*** | + | + | + | - | + | + |  |
| ***cnr2*** | + | + | - | - | + | + |  |
| ***ppara*** | + | + | + | + | + | + |  |
| ***pparg*** | + | + | + | + | + | + |  |
| ***gpr18*** | + | + | - | + | + | + |  |
| ***gpr55*** | + | - | - | - | + | + |  |
| ***gpr119*** | - | - | - | - | - | - |  |
| ***trpv1*** | + | + | + | + | + | + |  |
| ***trpv2*** | + | + | + | - | + | + |  |
| ***faah*** | + | + | + | + | + | + |  |
| ***naaa*** | + | + | + | + | + | + |  |
| ***htr1a*** | + | - | - | - | - | - |  |
| ***adora2A*** | + | + | + | + | + | + |  |
| ***adgrf1*** | + | + | + | + | + | + |  |

**Table-6:** Summary of the findings of the expression of the canonical and extended ECS in Rhesus macaques (*Macaca mulatta*).

| NHP | Colon | Heart | Kidney | Liver | MLN | Spleen | Visceral Fat |
| --- | --- | --- | --- | --- | --- | --- | --- |
| ***cnr1*** |  | - | - | - | - | + | + |
| ***cnr2*** |  | - | - | + | + | + | - |
| ***ppara*** |  | + | + | + | + | + | + |
| ***pparg*** |  | + | + | + | + | + | + |
| ***gpr18*** |  | - | - | - | - | + | - |
| ***gpr55*** |  | - | + | + | + | + | - |
| ***gpr119*** |  | - | - | - | - | + | - |
| ***trpv1*** |  | - | - | + | + | - | - |
| ***trpv2*** |  | - | - | + | + | + | + |
| ***faah*** |  | - | + | + | - | + | + |
| ***naaa*** |  | - | - | - | - | + | - |
| ***htr1a*** |  | - | + | + | - | + | - |
| ***adora2A*** |  | - | + | + | + | + | - |
| ***adgrf1*** |  | - | + | - | - | - | - |

### Both Canonical ECS Receptors Are Present In The Spleen Of Mice, Rats and NHP

The CB1R (*cnr1* gene) is most widely studied in the brain where it mediates antinociceptive effects, appetite regulation, and interacts with phytocannabinoids (20,21). However, CB1R is also present in multiple peripheral sites, including fat, lungs and reproductive organs where its plays roles in regulating inflammation and obesity (20,70). In contrast, CB2R is present primarily in peripheral organs, such as the spleen and MLN, and is most widely implicated in immune cell functions (71).

Here, we report that *cnr1* and *cnr2* mRNA were detected in peripheral organs of all three preclinical animal models, in agreement with existing studies. However, we expand on this knowledge to identify similarities and key differences among the model systems. *Cnr1* was most highly present in the visceral fat tissue for mice, while in rats its highest levels instead occurred in the kidney and colon. The NHP model had more limited *cnr1* where it was detectable only in the spleen and visceral fat. Similar findings occurred with *cnr2*. While *cnr2* was commonly detected in the MLN and spleen of all animal models, as expected, nuanced expression also existed where it was present in the rat colon, but not in the mouse. These findings denote key differences in *cnr1* mRNA not only between rodent and NHP models, but also between mice and rats.

Importantly, we determined that both canonical receptors were detected in the spleen of all three preclinical models, suggesting that it is a well conserved candidate to study the implications of CB1 and CB2 in health and disease. However, it must be acknowledged that we also identified important differences between mice, rats, and NHP. Indeed, we identified there was a striking absence of *cnr1* mRNA in the liver and of *cnr2* in the kidney for all three preclinical animal models, which is inconsistent with human expression patterns (20,72,73). This demonstrates an important limitation in translation across species. Further, this demonstrates the necessity for comparative ECS analyses to identify appropriate preclinical animal models to determine those that are best reflective of what occurs in humans.

### Nuclear Transcription Factors, *Ppara* and *Pparg,* Are Generally Well Conserved In All Organs Of Mice, Rats and NHP

PPARs are a group of ligand-activated nuclear hormone receptors (*ppara*, *pparb/d* and *pparg*) that interact with Retinoid X Receptor to act as transcription factors that regulate gene expression of genes involved in energy metabolism, glucose and fat metabolism, and inflammation (23,25,74,75). In humans, mice and rats, *ppara* is ubiquitous, but in rodents it has biased expression in energy requiring organs, including the heart, liver, and kidney (76–81). In contrast, PPARG is primarily present in the fat tissue in humans and mice, while rats have highest expression in the thymus (79–81).

Our PPARA and PPARG findings are in agreement with existing knowledge, except for their notable lack of detection in the spleen of mice. Indeed, except for this occurrence, PPARA and PPARG were the most highly conserved ECS receptor gene evaluated, having comparable detection among all organs across all preclinical models. Interestingly, *ppara* and *pparg* mRNA were detected ubiquitously in all evaluated organs in the NHP model. These findings demonstrate high translational potential for *ppara* and *pparg* and provide implications for evaluating how cannabinoids may impact energy homeostasis (74), macrophage activation, insulin sensitivity, (24,82,83), and anti-inflammatory pathways through NF-kB inhibition (24,84).

### Endocannabinoid-like GPRs Are Preferentially Expressed In Lymphoid Organs and The Visceral Fat

There is limited understanding of the endocannabinoid-like GPRs, which in humans is restricted to detection of *gpr18* and *gpr55* in lymphoid tissue and reproductive organs (85,86), and *gpr119* in the GI tract (87,88). Here we report that, overall, the GPR’s had limited expression across all preclinical models evaluated. Further, when detected, there were marked organ- and species-specific differences. The GPR’s were most consistently detected in the spleen, where *gpr18* and *gpr55* were expressed in all evaluated animal models. However, there was a noticeable lack of *gpr119* in the spleen, which was conserved among the mice, rat, and NHP models. This trend continued, as there was more widespread *gpr18 and gpr55* expression among all organs, although there were key species-specific differences. Indeed, *gpr18* was primarily restricted to the rat model (heart and spleen), which were not detectable in mice and NHP. In sum, these findings demonstrate that the rat model represents the best preclinical model to evaluate endocannabinoid-mediated GPR activation *in vivo.* Further, we identify the spleen as the most attractive therapeutic option to target the GPR’s as it has the most consistent expression across all evaluated models.

While their endogenous functions are incompletely understood, the GPR’s have clinically relevant implications, including *gpr18’s* roles in intracellular calcium, immunomodulation, cancer, metabolism and intraocular pressure (28,89–91); g*pr55’s* effects on bone physiology and intracellular signal transduction involving the activation of NF-κB, NFAT, CREB and ATF2 (29–31,91–94); and g*r119’s* involvement in glucose homeostasis and insulin secretion and sensitivity (103–107).

### TRPV1 and TRPV2 Nociception Channels Have Limited Translational Applicability to Humans

*Trpv1* and *trpv2* are ion channels that allow passage of essential ions (i.e., Na^2+^ and Ca^2+^) through the cell membrane (32,33). These ionotropic receptors are involved in noxious stimuli such as pain, heat, and inflammation and its expression is ubiquitous in humans (98,99). Here we report marked differences in these receptors. Rats had the most similar *trpv1* expression patterns to humans as it was widely expressed, whereas it was more limited in mice and NHP. While *trpv2* was more abundant in all three preclinical animal models, the only organ with shared expression among all three preclinical models was the spleen. Expression in the colon, heart and kidney was detected between rodents but not in NHP model. These discrepancies are profound in comparison to humans and demonstrates relatively poor translational potential. While these models are invaluable tools to evaluate the function of these receptors, care must be taken in drawing conclusions to the human condition. This suggests high potential for failure of preclinical endocannabinoid studies that aim to evaluate the roles of *trpv1* in hyperalgesia, body temperature control, diabetes, hormone secretion, epilepsy and hearing (98), as well as *trpv2* in cancer and cardiovascular dysfunction (99–102).

### FAAH and NAAA Endocannabinoid Metabolic Enzymes Are Ubiquitous In Rodents, But More Restricted In The NHP Model

FAAH and NAAA are important components of the ECS through endocannabinoid regulation that are ubiquitously expressed in humans (34,35,38). Our results identify that *faah* and *naaa* are ubiquitously expressed in the peripheral organs of rodents. While there were statistically significant differences among the organs, the mRNA for these metabolic enzymes were always detectable in mice and rats. Surprisingly, there was limited expression in the peripheral organs of the NHP model. Notably, *faah* was not present in the NHP heart and MLN, while *naaa* was undetectable in all organs except for spleen. This suggests that rodent models may have better preclinical utility to perform cannabinoid studies focused on targets of *faah* and *naaa*. This realization is important as these enzymes are essential in regulating endocannabinoid tone, which when dysregulation leads to pathology (103,104). Inhibiting these endocannabinoid catabolic enzymes is of major therapeutic interest as FAAH inhibitors are suggested as therapeutic targets for a group of diseases related to endocannabinoid level deficiencies termed “Clinical Endocannabinoid Deficiency Syndrome”, which have implications for migraine, fibromyalgia, and irritable bowel syndrome (103,104).

### *Htr1a*, *Adora2a* and *Adgrf1* are Poorly Conserved Among Mice, Rats, and NHP

*Htr1a* and *adora2a* have been primarily studied in the context of the brain, while *adgrf1* is known to be expressed in the kidney (39,40,42,105,106). We identified minimal conservation of *htr1a*, *adora2a*, and *adgrf1* among the three animal models. Our results corroborated that *htr1a* was minimally present in the periphery of both mice and rats. However, we were surprised to learn of its more widespread presence in the NHP model where it was detected partially in the kidney, liver, and spleen. In contrast, *adora2a* was present in the kidney and spleen for all three preclinical animal models. Even with these similarities, we observed marked species differences as *adora2a* was not present in the liver of mice, while it was expressed in the liver of rats and NHP. Similar trends occurred for adora2a in the heart and visceral fat as they were detected only in the rodent models. While rats had detectable a*dgrf1* in all organs tested, it was not present in the heart or spleen of mice. Interestingly, NHP had the most limited *adgrf1* expression as it was restricted to the kidney, presenting a limiting factor in clinical translation from rodents to NHP and therefore to humans.

Strikingly, there was little overlap among *htr1a*, *adora2a* and *adgrf1* in the animal models. In fact, each species had only one organ where these genes were co-expressed: visceral fat for mice, colon for rat, and kidney for NHP. This dissimilarity in expression patterns warns that caution must be taken when evaluating cannabinoid-mediated effects on *htr1a*, *adora2a*, and *adgrf1* in efforts to identify new therapeutic targets, as the potential for limited translation is high. This is the most poignant demonstration of the care that must be taken when selecting preclinical animal models for endocannabinoid studies. The translational limitations of these receptors has clinical implications as *htr1a* has been extensively studied as a target for mood disorders, a*dora2a* is suggested as a therapeutic target for neurodegenerative disorders, blood brain barrier integrity, immunosuppression, cancer, and angiogenesis (41,42), and a*dgrf1* is proposed as a novel therapeutic for cancer and inflammation (43,44,107).

## Conclusions

The endocannabinoid system is an incredibly attractive therapeutic target for many disorders where phytocannabinoids and receptor agonists/antagonists are being considered as novel treatment strategies. However, our findings demonstrate profound species- and organ-specific effects where there is limited overlap in expression pattern among mice, rat, and rhesus macaque preclinical models. We recommend that cannabinoid studies carefully consider the preclinical model to be included with respect to animal species, strain, genetic background, and even the site of procurement as there are reported variations in the same strains of rats obtained from different vendors (12). Further, cannabinoid formulation, vehicle in which it is reconstituted, metabolism, transport, mechanism of action, and their complex pharmacology should be considered to determine the right dose, formulation, and route of administration (12,108). Additionally, we urge scientists in the cannabinoid field to consider studying the relation between formulation, dosage, route of administration, diet, fed state, water availability, and cannabinoid pharmacokinetics. We also suggest considering age as an important factor as expression of endocannabinoid receptors, ligands and enzymes changes throughout the life course (68,69). It must also be considered that rodents are nocturnal animals, and their circadian rhythm may also impact results that can lead to discrepant findings with NHP and human preclinical studies. Finally, geographical location, time of administration, and time from administration to the actual experiment is performed should be taken into consideration. We anticipate our findings will provide insight for more rigorous experimental design of cannabis and cannabinoid translational research involving these preclinical animal models. Further, we hope that our findings will demonstrate the need to consider the extended ECS receptors that are abundantly expressed, activated by both endo- and phytocannabinoids, and that represent underlying mechanisms of action for these important lipid ligands.

## Data availability

The data gathered in this study is compiled and stored according to NIH data management, storing and sharing policies. Data will be available one year after the study has been published and can be accessed using the following link doi:10.17632/t6yd6j6bm6.1

## Acknowledgements

The authors would like to extend their gratitude towards the Retrovirus Laboratory at Johns Hopkins for their support and provision of samples from the historical rhesus macaque NHP samples included in this study, as well as Dr. Janice Clements for providing the funding that supported procurement and goals of the original research study in which these historical animals were involved. We would also like to acknowledge the veterinary staff that handled the rats and rhesus macaque at Johns Hopkins Bayview Campus and East Baltimore Campus, respectively. Research reported in this publication was supported by the National Institute on Drug Abuse of the National Institutes of Health under award number R01DA052859 and U01DA058527 (DWW) and the National Institute of Neurologic Disorders and Stroke award number K00NS118713 (CJW). The authors also acknowledge mentorship to DWW and procurement of pilot funds to ALE from the Johns Hopkins University Center for AIDS Research (P30AI094189), Diversity Supplement funded by R01-DA052859-03S1 to JJRF. REW is a Solomon H. Snyder Fellow at the Neuroscience Training Program at Johns Hopkins University. The content is solely the responsibility of the authors and does not necessarily represent the official views of the National Institutes of Health.

## Author’s contribution

JJRF participated in tissue collection, RNA extraction, cDNA synthesis, performed the experiments, determined the relative expression of most of the genes in all animal models, created all the graphs and data analysis and wrote the first draft of the manuscript. ALE participated in the mice necropsies and determined relative expression of the NHP genes *cnr1, gpr55, trpv1, trpv2*, and participated in editing the manuscript. CFM & EMW provided the rats included in this study, participated in necropsies and experimental feedback, and edited the manuscript. CJW & ASP provided the mice included in this project, participated on necropsies, and provided experimental feedback and edited the manuscript. REW performed experiments of mice *ppara*. DWW conceived the project idea and experimental design, provided direct guidance, funding for these experiments, and edited the manuscript.

